# Using DNase Hi-C techniques to map global and local three-dimensional genome architecture at high resolution

**DOI:** 10.1101/184846

**Authors:** Wenxiu Ma, Ferhat Ay, Choli Lee, Gunhan Gulsoy, Xinxian Deng, Savannah Cook, Jennifer Hesson, Christopher Cavanaugh, Carol B. Ware, Anton Krumm, Jay Shendure, C. Anthony Blau, Christine M. Disteche, William S. Noble, ZhiJun Duan

**Affiliations:** Department of Genome Sciences, University of Washington, USA; Department of Pathology, University of Washington, USA; Institute for Stem Cell and Regenerative Medicine, University of Washington, USA; Department of Comparative Medicine, University of Washington, USA; Department of Radiation Oncology, University of Washington, USA; Division of Hematology, Department of Medicine, University of Washington, USA; Howard Hughes Medical Institute Seattle, WA 98195-8056, USA; Present address: Department of Statistics, University of California, Riverside, USA; Present address: La Jolla Institute for Allergy and Immunology, USA

**Author notes:** Correspondence should be addressed at or.

**Keywords:** Chromatin, Chromosome, Chromosome conformation capture (3C), Hi-C, DNase Hi-C, Threedimensional (3D) genome architecture

## Abstract

The folding and three-dimensional (3D) organization of chromatin in the nucleus critically impacts genome function. The past decade has witnessed rapid advances in genomic tools for delineating 3D genome architecture. Among them, chromosome conformation capture (3C)-based methods such as Hi-C are the most widely used techniques for mapping chromatin interactions. However, traditional Hi-C protocols rely on restriction enzymes (REs) to fragment chromatin and are therefore limited in resolution. We recently developed DNase Hi-C for mapping 3D genome organization, which uses DNase I for chromatin fragmentation. DNase Hi-C overcomes RE-related limitations associated with traditional Hi-C methods, leading to improved methodological resolution. Furthermore, combining this method with DNA capture technology provides a high-throughput approach (targeted DNase Hi-C) that allows for mapping fine-scale chromatin architecture at exceptionally high resolution. Hence, targeted DNase Hi-C will be valuable for delineating the physical landscapes of *cis*-regulatory networks that control gene expression and for characterizing phenotype-associated chromatin 3D signatures. Here, we provide a detailed description of method design and step-by-step working protocols for these two methods.

**Highlights:**

- DNase Hi-C, a method for comprehensive mapping of chromatin contacts on a whole-genome scale, is based on random chromatin fragmentation by DNase I digestion instead of sequence-specific restriction enzyme (RE) digestion.
- Targeted DNase Hi-C, which combines DNase Hi-C with DNA capture technology, is a high-throughput method for mapping fine-scale chromatin architecture of genomic loci of interest at a resolution comparable to that of genomic annotations of functional elements.
- DNase Hi-C and targeted DNase Hi-C provide the first high-throughput way to overcome the RE-digestion-associated resolution limit of 3C-based methods.
- Step-by-step whole-genome and targeted DNase Hi-C protocols for mapping global and local 3D genome architecture, respectively, are described.

## 1. Introduction

The spatial organization of the genome has critical impacts on the various DNA-templated nuclear processes (e.g., transcription, DNA replication and repair) as well as on human development and disease [1–4]. Over the past decade, advances in technologies, especially the development of chromosome conformation capture (3C)-based high-throughput techniques (e.g., Hi-C), have yielded remarkable new insights into the principles of the three-dimensional (3D) genome organization [1, 2, 4–15]. It is increasingly clear that eukaryotic genomes are hierarchically organized in the nuclei [16–18]. Several conformational features corresponding to the distinct levels of genome organization have been identified, including whole chromosome territories [19], large-scale active and repressed compartments (A/B compartments) [20], specific domains, for example, topologically associated domains (TADs) [21, 22], lamin associated domains (LADs) [23, 24] and nucleolus associated domains (NADs) [25, 26], and ultimately, individual looping interactions between specific loci [27]. The detection of these chromatin conformation signatures associated with di erent scales of 3D genome organization require different levels of resolution. For example, chromosome territories and A/B compartments can be identified at the megabase scale, whereas TADs, LADs and NADs are at the sub-megabase scale. However, mapping of other fine-scale chromatin signatures, such as chromatin loops between *cis*-regulatory elements, requires much higher resolution [27].

The regulation of eukaryotic gene expression depends on transcription factors and cofactors acting on functional DNA elements such as promoters, enhancers, silencers and insulators, which are situated in the 3D nuclear space [1, 28]. Precisely mapping the spatial organization of *cis*-regulatory elements in the nucleus will lead to new insights into the principles underlying long-range gene regulation and how regulatory elements coordinate to achieve temporal and cell type-specific transcriptional regulation. Recent advances in genomic technologies have led to the systematic identification and annotation of functional DNA elements in mammalian genomes, including the human genome [29, 30]. The human genome contains many thousands of *cis*-regulatory elements (e.g., enhancers, silencers and insulators) that, in any given human cell, are located both proximal to and far away from the genes they control [31–34]. Although the precise mechanisms for gene regulation by distant-acting *cis*-regulatory elements remain largely elusive, it has been shown that chromatin looping-mediated physical contacts between gene promoters and other *cis*-regulatory elements are essential for transcriptional activities of the human genome [31–34].

ChIA-PET [35] and 3C derivatives such as 4C [36, 37], 5C [38], Hi-C [20], Capture-C [39], Capture Hi-C [40] and HiChIP/PLAC-seq [41, 42] can be used to map fine-scale chromatin architecture such as the chromatin interaction landscapes of *cis*-regulatory elements. However, since these 3C-based methods all rely on restriction enzyme (RE)-digestion to fragment chromatin, the ultimate resolution of fine-scale chromatin architecture mapped by these 3C-based methods is limited by the local distribution of RE sites, which is uneven across the genome (Fig. 1). Ultimately, a chromatin interaction identified by RE digestion-based 3C methods can only be described as the interaction between corresponding RE fragment pairs, and thus may not necessarily refer to the precise genomic location (up to single base pair resolution) of *cis*-regulatory elements [43], where the physical contact occurs.

**Fig. 1.**
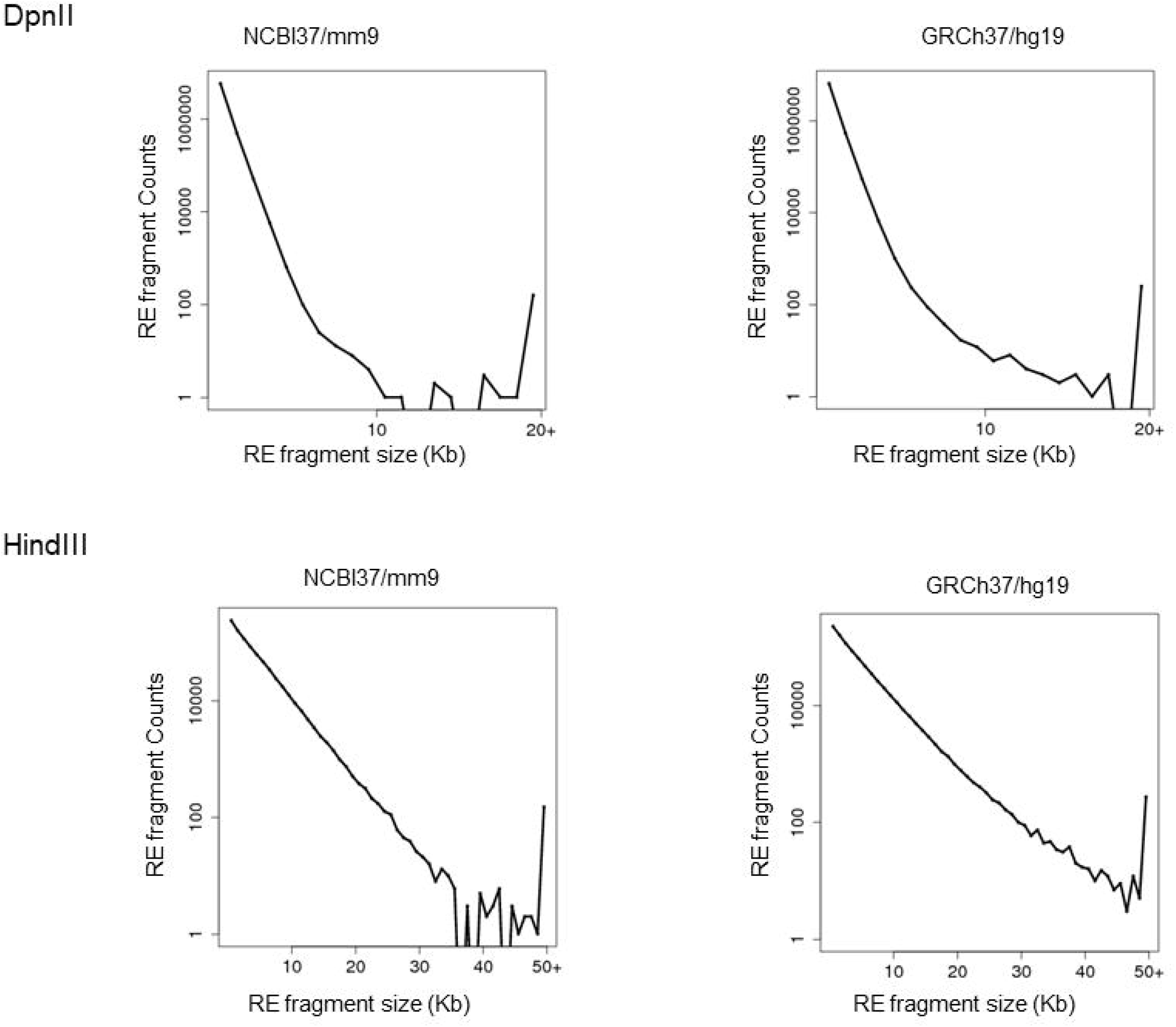
Size distribution of the projected restriction fragments in mouse and human genomes after the *in silico* digestion with DpnII and HindIII, two REs frequently used in Hi-C assays. Note that for the 4 bp-cutter DpnII, about 8.73% (559,888 of 6,415,145) of fragments (RE-fragment counts) generated in the mouse genome (NCBI/mm9) and 8.58% (611,434 of 7,127,561) in the human genome (GRCH37/hg19) are larger than 1 kb. Importantly, the DpnII fragments with a size > 1kb cover about 31.25% of the mouse genome and 33.28% of the human genome, respectively. For the 6 bp-cutter HindIII, about 20.38% (167,820 of 823,331) of fragments generated in the mouse genome and 22.74% (190,456 of 837,599) in the human genome are larger than 5 kb. Notably, the HindIII fragments with a size > 5 kb cover about 52.22% of the mouse genome and 58.33% of the human genome, respectively.

To overcome the resolution limitation of RE-based 3C methods, and toward the development of a single-base-resolution method for mapping chromatin interactions in the future, we recently developed DNase Hi-C, a method for comprehensively mapping chromatin interactions on a whole-genome scale in living cells [44] (Fig. 2). Although, like all the traditional 3C-based methods, the core concept of DNase Hi-C is also based on proximity ligation, DNase Hi-C employs DNase I instead of an RE to fragment the crosslinked chromatin (Fig. 2). Importantly, the sequence-independent digestion of chromatin by DNase I enables DNase Hi-C to overcome the resolution limitation associated with RE-based 3C methods. Subsequently, we have combined DNase Hi-C with DNA capture technologies to develop a method for fine-mapping chromatin architecture of specific genomic regions of interest at unprecedented resolution in a massively parallel fashion, termed “targeted” DNase Hi-C [44] (Fig. 2). We have applied DNase Hi-C to a diversity of biological systems, including yeast cells [45], mouse cell lines and tissues [46], and human cell lines [44, 47]. We have also developed an *in situ* version of DNase Hi-C, termed *in situ* DNase Hi-C [46–48].

**Fig. 2.**
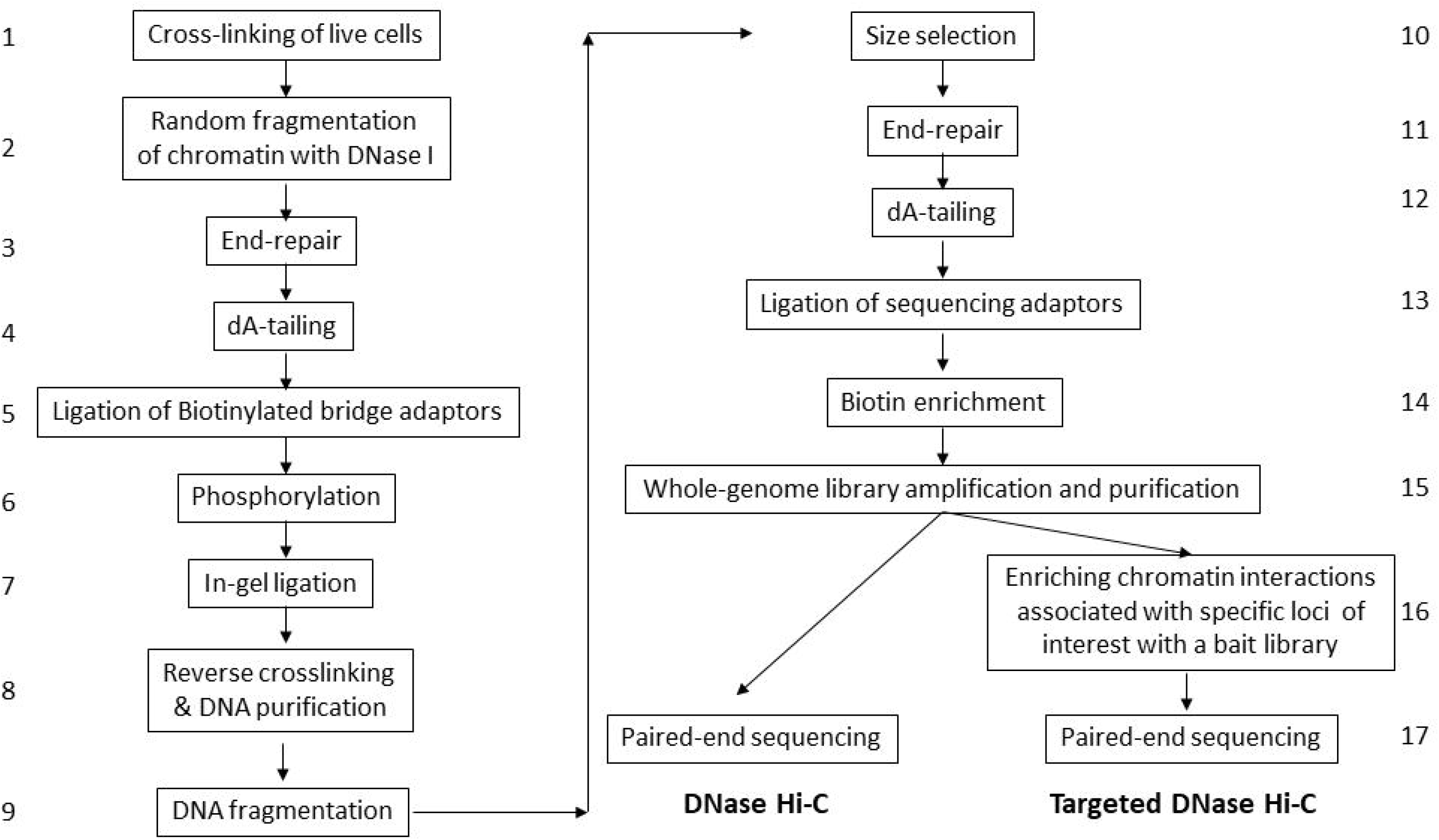
The workflow of whole-genome and targeted DNase Hi-C assays.

## 2. Overview of DNase Hi-C and targeted DNase Hi-C

The use of DNase I digestion leads to many differences between the protocol of DNase Hi-C and that of conventional RE-based Hi-C (Fig.2). Briefly, cells are cross-linked with formaldehyde; chromatin is then randomly fragmented by DNase I. The resulting chromatin fragments are end-repaired and dA-tailed, then marked with a biotinylated internal adaptor. Proximity ligation is carried out by burying the biotin-labeled fragments inside solidified agarose gel to favor ligation events between the cross-linked DNA fragments. The resulting DNA sample contains ligation products consisting of chimeric DNA fragments that were originally in close spatial proximity in the nucleus, marked with biotin at the junction. A whole-genome chromatin interaction library is created by shearing the DNA and selecting the biotin-containing fragments with streptavidin magnetic beads. After linear library amplification, a DNase Hi-C library is generated and can be sequenced to identify genome-wide chromatin contacts. Alternatively, the chromatin contacts associated with a specific group of loci of interest are then enriched by a bait library targeting the loci via in-solution hybridization-based DNA capture (targeted DNase Hi-C). Finally, quantitation of the enriched chromatin interactions is achieved through massively parallel deep sequencing.

### 2.1. Formaldehyde-mediated cell crosslinking and cell lysis

The highly dynamic nature of chromosome higher order structure leads to cell-to-cell variation of chromatin interactions within any given cell population, even if the cells of the population appear to be functionally homogeneous. Hence, at any given time, chromatin interactions occurring in individual cells of a cell population likely differ and not all chromatin interactions, especially infrequent interactions, occur in all the cells. Consequently, starting the protocol with a larger number of cells leads to higher complexity of the DNase Hi-C library and higher achievable resolution. Therefore, although a DNase Hi-C assay can be carried out with as low as a few thousand cells, which is suitable for rare cell populations, we recommend starting with 1-2 million cells for a whole-genome DNase Hi-C assay and 3-5 million cells for a targeted DNase Hi-C assay (or even more cells, depending on the size of the targets and the desired resolution).

DNase Hi-C assays can be carried out in various cell types, such as yeast cells, plant cells, cultured mammalian cells and tissues. To capture chromatin interactions, all types of cells can be crosslinked with formaldehyde, which is the most widely used chemical crosslinking agent in chromatin immunoprecipitation (ChIP) assays and in 3C-based methods, due to several advantageous features compared to other crosslinking agents [49]. However, different types of starting materials may require different conditions for preparing crosslinked cells or nuclei. For example, to conduct a DNase Hi-C assay with yeast cells, the cells need to be cross-linked with 1% (by volume) formaldehyde for 10 min at room temperature, then the yeast cell wall is removed and spheroplasts are isolated [49]. To carry out DNase Hi-C in tissue cells, tissues need to be first homogenized, for example, with a Dounce tissue grinder, and the concentration of formaldehyde used for crosslinking may be tissue-specific [46]. Even for mammalian cell cultures, different cell types may also require different crosslinking conditions. For example, for suspension cells, such as K562, Jurkat, and lymphoblastoid cells, crosslinking can be carried out with 1% formaldehyde for 10 min at room temperature, the so-called standard crosslinking condition. Similarly, for monolayer adherent cell cultures, such as Hela, IMR90 and HEK293 cells, fixation can be performed under standard conditions without detaching cells from their growth surface. However, for mouse and human embryonic stem cells (ESCs), because they usually aggregate together and form big clumps during growth, efficient fixation of these cells requires one of two alternative approaches: preparing single-cell suspension before carrying out crosslinking under the standard condition, or crosslinking the cells without detaching them from their growth surface with increased formaldehyde concentration or longer fixation time. Because the 3D genome architecture can be disturbed during the preparation of single-cell suspensions, the latter strategy is preferred. Finally, because serum is enriched in proteins and can therefore interfere with formaldehyde-mediated crosslinking, it is recommended to carry out crosslinking with serum-free media. Fixed cells are then treated sequentially with hypotonic buffer containing IGEPAL CA-630 and SDS to open the chromatin for the following enzymatic reactions, while retaining the 3D chromatin conformation.

### 2.2. Fragmentation of crosslinked chromatin by DNase I digestion

Our protocol is based on a sequence-independent fragmentation of cross-linked chromatin by DNase I, which replaces the sequence-specific RE digestion used in traditional 3C methods. It is well known that DNase I, an endonuclease, nonspecifically cleaves DNA to release oligonucleotide products with 5’-phosphorylated and 3’-hydroxylated ends. We found that DNase I digestion of cross-linked chromatin is robust and controllable by adjusting the enzyme concentration and reaction temperature (Fig. 3A). Furthermore, digestion of crosslinked chromatin by DNase I into small DNA fragments (with a size less than 1 kb) does not lead to a significant loss of DNA, and the sequence reads of a DNase I Hi-C library cover a higher percentage of the mappable regions of the human genome than that of a typical Hi-C library [44]. Hence, we have demonstrated that DNase I digestion is an appropriate choice for the purpose of comprehensively mapping chromosome conformation signatures.

**Fig. 3.**
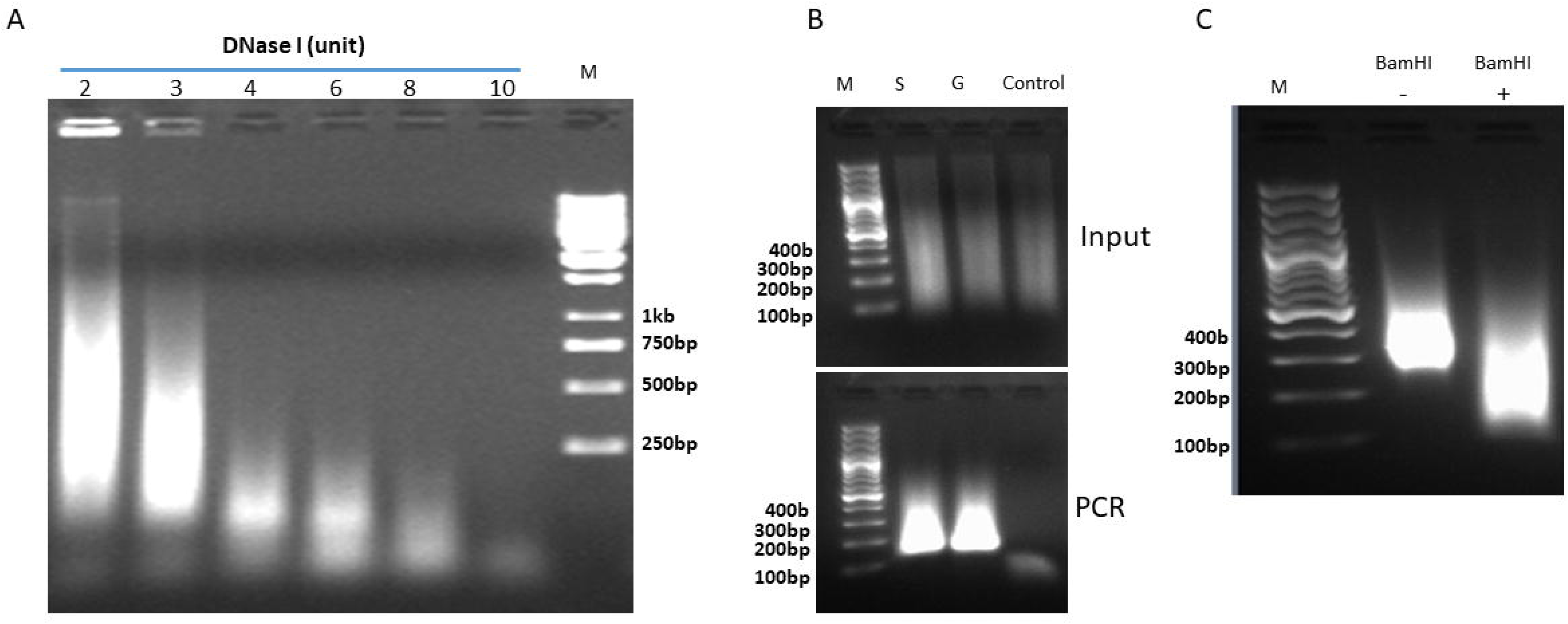
Quality control (QC) of experimental steps in the DNase Hi-C protocol. (A) Example showing titration of DNase I digestion of crosslinked chromatin (see protocol step 25). About 5 ⨯ 10^6^ crosslinked K562 cells (fixed in 1% formaldehyde) were lysed and divided into 6 equal aliquots, and each aliquot was digested with 2-10 units of DNase I at RT for 5 min. Genomic DNA was then isolated and 1/10 of the purified genomic DNA from each sample was subjected to gel electrophoresis in a 2% agarose gel. M, DNA size ladder. (B) Example showing QC of the enzymatic reactions in the DNase Hi-C protocol. About 2 ⨯ 10^6^ crosslinked K562 cells (fixed in 1% formaldehyde) were subjected to cell lysis, DNase I digestion, end-repair, and dA-tailing according to the DNase Hi-C protocol. After dA-tailing, the sample was divided into 3 equal aliquots, and ligated to the Illumina Y adaptor in solution (S), in a 0.4% low melting gel (G), or not ligated as a control (Control). Upper panel, about 300 ng purified genomic DNA from each sample (Input) was resolved in a 2% agarose gel; Lower panel, the corresponding PCR products of each input (1 ng DNA template) amplified using Illumina PCR primers (11 PCR cycles). M, 100 bp DNA ladder. (C) Example of the BamHI QC experiment (see protocol steps 135-137) performed on a K562 whole-genome DNase Hi-C library. Electrophoresis was carried out on a 2% DNA agarose gel. M, 100 bp DNA ladder.

### 2.3. Immobilization of nuclei/chromatin complexes with SPRI magnetic (AMPure) beads

SPRI (Solid Phase Reversible Immobilization) Magnetic Beads (e.g., Ampure XP) have been widely used for DNA and RNA purification. We found that, in addition to DNA and RNA, these beads can bind to intact nuclei and chromatin complexes. Hence, in the protocols of DNase Hi-C and *in situ* DNase Hi-C, SPRI beads are employed as carriers to immobilize nuclei/chromatin complexes. This strategy enables efficient removal of DNase I and of the DNA modifying enzymes used in the various enzymatic reactions of the protocols, including DNA polymerase I, large (Klenow) fragment, T4 DNA polymerase, and Klenow fragment (3’–>5’ exo-). SPRI beads are also essential for the efficient removal of free, unligated internal bridge adaptors following the experimental step of bridge-adaptor ligation, as well as the removal of low-molecular-weight DNA fragments that might escape the nucleus following cell lysis and chromatin digestion. Also, when starting with a low number of cells (e.g., less than 1 million cells), the SPRI beads (usually with a brown color) can help to visualize the cell/nuclei pellets throughout the protocol.

### 2.4. Marking chromatin ends with biotin

Marking the juxtaposition of two interacting genomic locations for its efficient enrichment in a Hi-C library usually requires labelling the chromatin ends with biotin. In the DNase Hi-C protocol, the biotin marking of chromatin fragments is achieved by ligation of the biotinylated-bridge adaptors through T-A ligation. Because DNase I digestion of chromatin generates a heterogeneous mixture of fragment ends composed of 5’– and 3’-overhangs of varying lengths as well as blunt ends, prior to ligation of the bridge adaptors, it is necessary to repair these ends with the DNA modifying enzymes Klenow fragment and T4 DNA polymerase, and to create dA-tailed ends with the enzyme Klenow fragment (3’–>5’ exo-).

### 2.5. In gel proximity ligation

Embedding large-sized DNA fragments in agarose gels to prevent random degradation and subsequently carrying out enzymatic reactions in gel has been previously used in several molecular methods such as DNase-seq [50]. In DNase Hi-C, the chromatin fragments generated by DNase I digestion are generally much smaller than those resulting from RE digestion. The large number and small size of the DNase fragments might increase the chance of random inter-molecular collisions during proximity ligation. Hence, to reduce the background noise arising from random intermolecular ligations between DNA fragments that are not cross-linked to each other, proximity ligation in the DNase Hi-C protocol is carried out by immobilizing nuclei/chromatin complexes into a low-melting agarose gel. Using low-melting agarose with its melting and gelling temperatures of 65.5°C and 25°C, respectively, allows the molten solution to be equilibrated to 37°C before mixing with the nuclei/chromatin complexes to help preserve nuclei and chromosome structure. Also, the concentration of the gel is crucial to achieve the best effect. On the one hand, the higher the gel concentration, the lower the mobility of the embedded nuclei/chromatin complexes. On the other hand, higher gel concentration will reduce the accessibility of DNA fragments by T4 DNA ligase and lead to lower ligation efficiency. We found the optimal gel concentration to be 0.4%.

### 2.6. Enriching a subset of chromatin interactions of interest from whole-genome DNase Hi-C libraries by using DNA capture technologies (targeted DNase Hi-C)

Chromatin interactions associated with specific genomic regions of interest can be accurately enriched from a DNase Hi-C library without incurring severe biases by using DNA capture technology. These methods have been widely used in targeted genome sequencing, such as exome sequencing [51]. However, there is a significant difference between capturing chromatin interactions from a DNase Hi-C library and capturing DNA sequences from a simple genomic DNA library. In exome capture assays, each DNA fragment of a target region is fully covered by the designed hybridization probes, provided that its sequence is uniquely mappable. In contrast, in a DNase Hi-C library, each valid DNA fragment is chimeric, consisting of two interacting fragments linked via the internal adaptor (Fig.4A). Probes can only be designed to cover the portion of the fragment that is located within the target region of interest but not the other portion of the fragment, which is unknown and to be identified (Fig.4A). Nevertheless, we found that the efficiency of capturing chromatin interactions from a DNase Hi-C library is similar to that of capturing single DNA fragments from a genomic library [44], indicating that targeted DNase Hi-C can be used to efficiently capture subsets of chromatin interactions in a genome.

**Fig.4.**
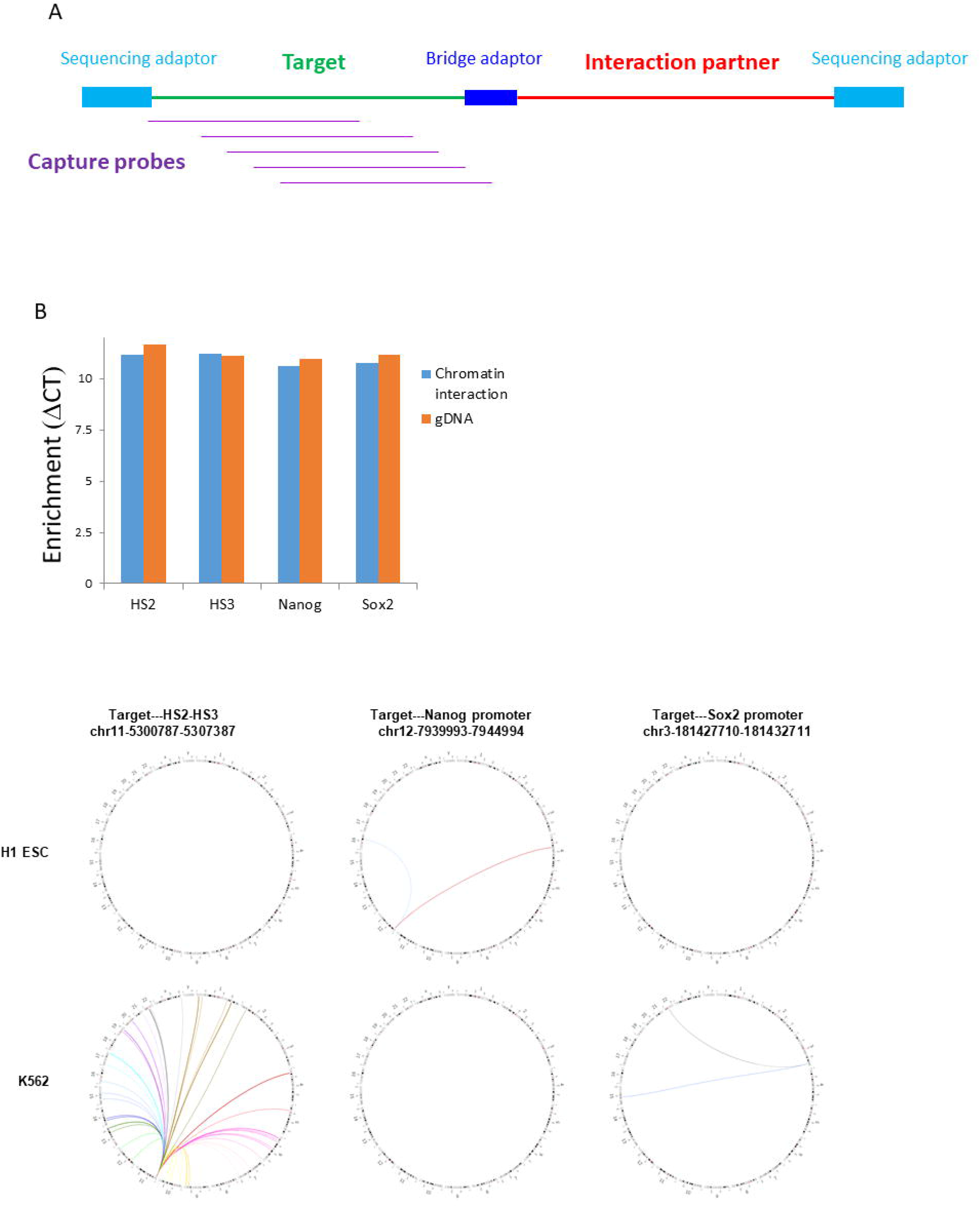
Enrichment of a subset of chromatin interactions of interest from whole-genome DNase Hi-C libraries by DNA capture. (A) Cartoon to illustrate capture of target-associated chromatin interactions using probes designed against a specific target. The structure of the chimeric DNA fragments (templates) in DNase Hi-C libraries is shown, including sequencing adaptors and bridge adapter. (B) Example of using real-time PCR to assess enrichment efficiency of target-associated chromatin interactions in targeted DNase Hi-C assays. Enrichment is comparable between a whole-genome DNase Hi-C library (chromatin interaction) and a control genomic library (gDNA) at four loci (HS2, HS3, Nanog, Sox2) using real time PCR.

Due to their simplicity, scalability, high efficiency and high reproducibility, in-solution hybridization-based methods are most commonly used, among all the presently available DNA capture technologies. In the present version of the targeted DNase Hi-C protocol the Roche NimbleGen SeqCap EZ technology has been chosen among the commercially available insolution hybridization platforms, to capture chromatin interactions from DNase Hi-C libraries, for two reasons. First, the size of its DNA probes is flexible (55-105 nt), which is well-suited for enriching the chimeric ligation products in a DNase Hi-C library. While the size of each chimeric DNA fragment is defined by size fractionation, the size of each portion of the chimeric DNA fragments is widely variable. Second, the SeqCap design algorithm is more advanced than that offered by other companies and produces the highest probe density for any given target region. However, the Agilent Sureselect technology may have its own advantage by using RNA probes, since the affinity of RNA:DNA hybridization is higher than that of DNA:DNA. It should be noted that targeted DNase Hi-C can, in principle, be adapted for other array-based or in-solution-based capture methods.

#### 2.6.1. Complexity of targeted DNase Hi-C libraries

Similar to the situation in exome-capture assays, we found that pre-capture and post-capture PCR amplification and the size of target regions are the three major factors that affect the complexity of targeted DNase Hi-C libraries. Hence, to maximize the complexity of targeted DNase Hi-C libraries, it is preferable to keep a low number of PCR cycles for both pre-capture PCR (for DNA template amplification) and post-capture PCR (for targeted DNase Hi-C library amplification). Accordingly, to obtain a sufficient amount of material for sequencing with reduced PCR amplification, one can pool together PCR products obtained from the same sample. Moreover, similar to traditional 3C methods, it might be important to increase the amount of starting material (e.g., to more than 5 million cells) in some cases to ensure sufficient complexity of targeted DNase Hi-C libraries.

#### 2.6.2. Efficiency of targeted DNase Hi-C

We found that the variability of captured sequence read pairs between different genomic targets in a targeted DNase Hi-C assay mainly results from two sources: (1) unevenness of chromatin interactions associated with each target in the template DNase Hi-C library, which is a similar problem to that of RE-based Hi-C libraries, and (2) unevenness of DNA capture efficiency. There two major factors affect the enrichment efficiency of targeted DNase Hi-C assays. First, similar to exome-capture assays, the structure of the chimeric DNA fragments in DNase Hi-C libraries (Fig 4A), i.e., the size of the genomic fragments and the length of the adaptors, can strongly influence the enrichment efficiency of targeted DNase Hi-C assays. Hence, to minimize repeat-mediated cross-hybridization, we reduce the size of the genomic fragments to 200-350 bp in our DNase Hi-C libraries by sonication. Similarly, to prevent adaptor-mediated cross-hybridization, we use short PCR primers to amplify the DNA templates for targeted DNase Hi-C capture in the pre-capture PCR so that the adaptors associated with the DNA templates are of minimal length.

Second, the genomic properties (e.g., GC content) of a target can severely affect its DNA capture enrichment efficiency. It has been observed that, in genome-capture assays, the enrichment of *cis*-regulatory noncoding regions is lower (~70%) than that of the consensus coding DNA sequence [52]. To achieve similar coverage, capturing *cis*-regulatory noncoding loci requires 20-40% more sequencing data than capturing coding sequences [33]. Hence, with respect to coverage uniformity, unevenness across genomic regions with various GC contents has been a general problem in whole genome sequencing and exome-capture assays. Similarly, although the enrichment efficiency of the majority of the targets is very similar in a targeted DNase Hi-C assay, we indeed have observed unevenness in capture efficiency among different target regions [44]. Owing to the high GC-content nature of the *cis*-regulatory elements (e.g., promoters and enhancers), when using targeted DNase Hi-C to enrich promoter– or enhancer-associated chromatin interactions, optimization in probe design (e.g., altering the probe length and increasing the density of probes) will be helpful for maximizing the enrichment efficiency.

#### 2.6.3. Fidelity of targeted DNase Hi-C

In addition to enrichment efficiency, the other most important success criterion of a targeted DNase Hi-C assay is whether it can accurately capture the chromatin interaction patterns occurring in a genome. Two observations have demonstrated the faithfulness of targeted DNase Hi-C. First, targeted DNase Hi-C exhibits the relationship between genomic distance and average interaction frequency expected of polymer chromatin fibers [44]. Second, we have observed a high correlation of sequence read pairs associated with each target between targeted DNase Hi-C and the parent DNase Hi-C libraries [44].

### 2.7. Quality control of DNase Hi-C and targeted DNase Hi-C libraries

During the construction of DNase Hi-C and targeted DNase Hi-C libraries, quality control (QC) analysis should be carried out at several critical steps.

#### 2.7.1. DNase I digestion efficiency

The DNase I digestion efficiency can be examined by DNA gel electrophoresis (Fig. 3). Genomic DNA can be isolated from a small portion (~5%) of DNase I digested sample, and the size distribution of the DNA fragments checked by running a 1% agarose gel or a precast 6% polyacrylamide DNA gel. In general, an efficient DNase I digestion will result in the majority of DNA fragments with a size less than 1 kb (Fig. 3). We recommend this QC step when applying DNase Hi-C protocol to a new cell type.

#### 2.7.2. Efficiency of the enzymatic reactions (end-repair, dA-tailing, and adaptor ligation reactions) for biotin-labeling chromatin ends

The sensitivity of DNase Hi-C assays is largely determined by the efficiency of the three consecutive enzymatic reactions, Klenow enzyme-catalyzed DNA end-repair, Klenow (Exo^-^)-mediated dA-tailing and ligation of the biotinylated-bridge adaptors to the dA-tailed chromatin ends by T4 DNA ligase. To examine the efficiency of these reactions, a small portion (~5-10%) of dA-tailed sample can be taken to carry out a control ligation in parallel to the experimental ligation of the biotinylated-bridge adaptors. In the control ligation, the dT-tailed Illumina sequencing Y-adaptors are ligated to the dA-tailed chromatin ends. Genomic DNA is then isolated and the ligation efficiency examined by performing quantitative PCR (qPCR) with the appropriate Illumina PCR primers (Fig. 3B). If amplification signal appears before 10 PCR cycles using 10 ng of genomic DNA as a template, it suggests that the efficiency of the upstream end-repair and dA-tailing steps are acceptable. We recommend this QC step when applying the DNase Hi-C protocol to a new cell type.

#### 2.7.3. BamHI digestion analysis of whole-genome and targeted DNase Hi-C libraries

The biotinylated bridge adaptor for marking the chromatin ends in the DNase Hi-C protocol is designed in such a way that a BamHI enzyme site will be formed at the junction when two chromatin fragments ligate with each other through the bridge adaptors during the in-gel proximity ligation. Hence, in whole-genome or targeted DNase Hi-C libraries, each valid DNA fragment is chimeric, consisting of two interacting genomic fragments via the bridge adaptor with a BamHI site at the joint. Therefore, DNase Hi-C libraries can be validated by carrying out a BamHI digestion assay, i.e., for a valid DNase Hi-C library, there should be an apparent size shift of DNA fragments after BamHI digestion (Fig. 3C). We recommend applying this QC step to every whole-genome or targeted DNase Hi-C library prior to sequencing.

#### 2.7.4. Estimation of enrichment efficiency of a targeted DNase Hi-C library using qPCR

The enrichment efficiency of a targeted DNase Hi-C library can be estimated using qPCR before the library is subjected to sequencing analysis (Fig. 4B). PCR primers to specific target regions can be designed and qPCR carried out by using the targeted DNase Hi-C library or the parent whole-genome DNase Hi-C library as templates. The enrichment of a specific target can be assessed by comparing its amplification rate from the target DNase Hi-C library with that from the parent whole-genome DNase Hi-C library (Fig.4B). Since the capture efficiency of a specific target in a targeted DNase Hi-C assay is determined by multiple factors and is likely to be uneven in enrichment between individual targets, as discussed in Section 2.6, we recommend examining enrichment at multiple individual targets simultaneously using this qPCR assay to ensure the accuracy of the assessment.

### 2.8. Sequencing

Whole-genome and targeted DNase Hi-C libraries are sequenced using Illumina 75 bp, 100 bp or 150 bp paired-end sequencing. In general, longer paired-end reads can increase mapping efficiency. Moreover, longer reads (e.g., 150 bp) are required for efficiently identifying allele-specific chromatin interactions based on single nucleotide polymorphisms (SNPs) [46, 48].

## 3. Detailed protocols

### 3.1. Crosslinking of cells

As discussed in section 2.1, different starting materials may require different conditions for preparing crosslinked cells. In general, 1-2 ⨯ 10^6^ cells are sufficient for generating a whole-genome DNase Hi-C library. However, we suggest 3-5 ⨯ 10^6^ cells for a targeted DNase Hi-C assay and growing more cells for cross-linking to provide technical replicates if necessary. Below are the working protocols for adherent monolayer and suspension cells, respectively.

#### 3.1.1. Adherent monolayer cells

1. Aspirate the medium and add 10 ml of fresh medium without serum per 10 cm-plate.
2. Crosslink the cells by adding 280 µl of 37% formaldehyde (1% final concentration). Mix gently, immediately after addition of formaldehyde.
3. Incubate at room temperature (RT) for 10 min.
4. Add 560 ul of 2.5 M glycine to quench the crosslinking reaction, mix well.
5. Incubate for 10 min at RT to stop cross-linking completely.
6. Discard the supernatant by aspiration and wash the crosslinked cells with 1 x PBS once.
7. Scrape the cells from the plates with a cell scraper and transfer to a microtube.
8. Split the crosslinked cell suspension into aliquots of 1-2 ⨯ 10^6^ cells (in 1.7 ml microtubes).
9. Centrifuge the cross-linked cells at 800*xg* for 3 min.
10. Cells can be snap-frozen in liquid nitrogen and stored at -80°C for at least 1.5 years or one can continue with cell lysis (section 3.2).

Note, for those adherent cells that grow in an aggregated state, such as mouse and human ESCs, higher concentration of formaldehyde may be necessary (e.g., 2.5% final concentration).

#### 3.1.2. Suspension cells

1. Gently pellet the cells (about 2-3 ⨯ 10^7^) by spinning at 800*xg* for 5 min at RT.
2. Discard the supernatant.
3. Thoroughly resuspend the pellet in 45 ml of fresh culture medium without serum. Break
cell clumps by pipetting up and down.
4. Crosslink the cells by adding 1.25 ml of 37% formaldehyde in one shot (1% final
concentration). Mix quickly by inverting the tube several times.
5. Incubate at RT for 10 min. Gently invert the tube every 1-2 min.
6. Add 2.5 ml of 2.5 M glycine to quench the cross-linking reaction, mix well.
7. Incubate for 10 min at RT to stop cross-linking completely.
8. Pellet the crosslinked cells at 800*xg* for 3 min.
9. Discard the supernatant by aspiration and resuspend the cells with 1 x PBS (1ml PBS per 10^6^ cells).
10. Split the crosslinked cell suspension into aliquots of 1-2 ⨯ 10^6^ cells (in 1.7 ml microtubes).
11. Centrifuge the cross-linked cells at 800*xg* for 3 min.
12. Discard the supernatant by aspiration.
13. Cells can be snap-frozen in liquid nitrogen and stored at -80°C for at least 1.5 years or one can continue with cell lysis (section 3.2).

### 3.2. Cell lysis and chromatin digestion with DNase I

14. Resuspend one crosslinked cell aliquot (1-2 ⨯ 10^6^ cells) in 1 ml of ice-cold cell lysis buffer containing protease inhibitor cocktail.
15. Incubate on ice for 10 min.
16. Centrifuge for 5 min at 2,500*xg* at RT.
17. Discard the supernatant and resuspend the pellet in 1 ml of 1x TE lysis buffer.
18. Incubate at 37^o^C for 10 min.
19. Centrifuge at 2,500*xg* at RT for 20 sec.
20. Resuspend the pellet in 400 µl of 0.5 x DNase I digestion buffer containing 100 units of RNase A, mix well.
21. Incubate at 37^o^C for 10 min.
22. Add appropriate amount (e.g., 3-6 units) of DNase I, mix well.
23. Incubate at RT for 5 min.
24. Add 80 µl of 5 x stop solution, mix well. Save 20µl of lysate for assessing the DNase I digestion efficiency if desired (see 25).
25. To assess DNase I digestion efficiency add 70 µl of 1x TE lysis buffer and 10 µl of Proteinase K (20 mg/ml) to the saved 20 µl of lysed cells from the previous step. Incubate for 30 min at 65°C. Purify DNA by the Qiaquick PCR purification kit. Check size distribution of the DNA fragments by running the sample on 1% agarose gel. In general, efficient DNase I digestion will result in the majority of DNA fragments with a size less than 1kb (Fig. 3A).
26. Add 1 ml of AMPure XP beads, mix well and distribute the cell-beads mixture between three 1.7 ml microtubes (about 500 µl per tube).
27. Incubate at RT for 5 min, and place the three tubes in a DynaMag-Spin magnet for 2 min.
28. Discard the supernatant and wash the beads twice with 1 ml of 80% ethanol and combine the beads of the three microtubes into one microtube. Briefly spin down the beads and remove the residual ethanol as completely as possible, then air dry the beads for no more than 2 min.
29. Resuspend immediately the beads with 100 µl of water, ready for the next end-repair step.

Note, the amount of DNase I used at step 22 can vary depending on the cell type being studied. Too much DNase I will result in over-digestion and loss of DNA, whereas too little DNase I will lead to inefficient chromatin fragmentation. When applying the protocol on a new cell type, we recommend carrying out a DNase I optimization experiment using varying amounts of DNase I, and then examining the size distribution pattern following digestion (Fig. 3), as described in step 25.

### 3.3. End repair

30. Prepare the End-Repair master mix as follows:
31. Add the100 µl End-Repair mastermix to the DNase I digested sample from step 29. Mix well by pipetting up and down.
32. Incubate at RT for 1 h.
33. Add 200 µl of 20% PEG buffer to the tube, mix thoroughly by pipetting up and down.
34. Incubate at RT for 5 min, and place the tube in a DynaMag-Spin magnet for 2 min.
35. Discard the supernatant and wash the beads twice with 1 ml of 80% ethanol. Briefly spin down the beads and remove the residual ethanol as completely as possible, then air dry the beads for no more than 2 min.
36. Resuspend immediately the beads in 100 µl of water, ready for the next dA-tailing step.

| Fill-in mix (add in this order) | Per reaction |
| --- | --- |
| Water | 70 μl |
| 10x T4 ligase buffer | 20 μl |
| 10 mM dNTPs | 5 μl |
| T4 DNA Polymerase (5U μl) | 2 μl |
| Klenow (5U/μl) | 3 μl |

### 3.4. dA-tailing

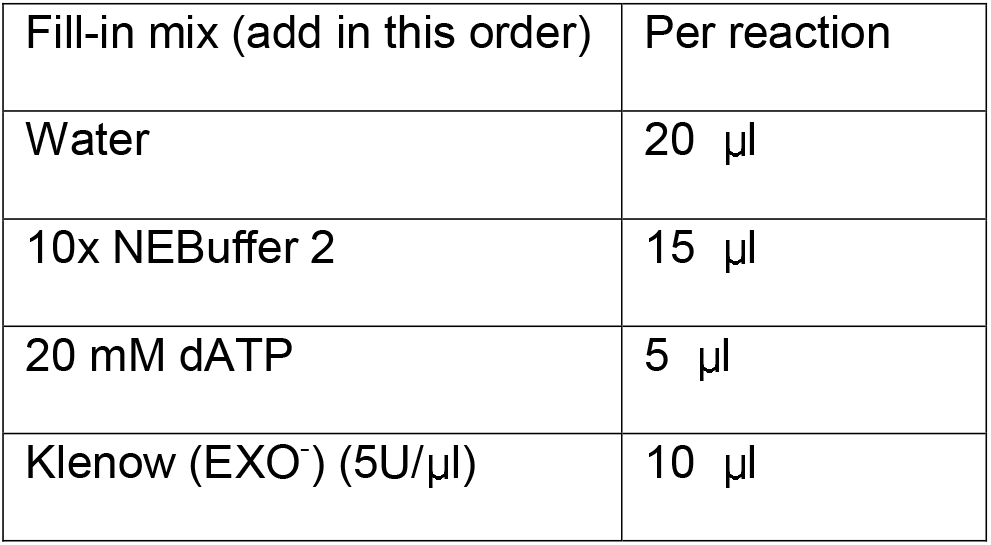
37. Prepare the dA-Tailing master mix as follows:
38. Add the 50 μl dA-Tailing master mix to the end-repaired sample from step 36. Mix well by pipetting up and down.
39. Incubate at 37^o^C for 1 h.
40. Add 150 µl of 20% PEG buffer to the tube, mix thoroughly by pipetting up and down.
41. Incubate at RT for 5 min, and place the three tubes in a DynaMag-Spin magnet for 2 min.
42. Discard the supernatant and wash the beads twice with 1 ml of 80% ethanol. Briefly spin down the beads and remove the residual ethanol as completely as possible, then air dry the beads for no more than 2 min.
43. Resuspend immediately the beads in 30 µl of water, ready for the next step

| Fill-in mix (add in this order) | Per reaction |
| --- | --- |
| Water | 20 μl |
| 10x NEBuffer 2 | 15 μl |
| 20 mM dATP | 5 μl |
| Klenow (EXO<sup>-</sup>) (5U/μl) | 10 μl |

### 3.5. Ligation of the Biotinylated Bridge adaptors to chromatin ends

44. Prepare the ligation master mix as follows:
45. Add the 70 µl ligation master mix to the dA-tailed sample from step 43. Mix well by pipetting up and down.
46. Incubate at 16^o^C overnight.
47. Add 5 µl of 10% SDS to the reaction.
48. Add 195 µl of ddH_2_O to bring up the volume to 300 µl, Mix thoroughly by pipetting up and down.
49. Add 300 µl of 20% PEG buffer to the tube, mix thoroughly by pipetting up and down.
50. Incubate at RT for 5 min, and place the tube in a DynaMag-Spin magnet for 2 min.
51. Discard the supernatant and wash the beads once with 1.5 ml of 80% ethanol.
52. Briefly spin down the beads and remove the residual ethanol as completely as possible.
53. Resuspend the beads in 200 µl of ddH_2_O, mix thoroughly by pipetting up and down.
54. Add 165 µl of 20% PEG buffer to the tube, mix thoroughly by pipetting up and down.
55. Incubate at RT for 5 min, and place the three tubes in a DynaMag-Spin magnet for 2 min.
56. Discard the supernatant and wash the beads twice with 1 ml of 80% ethanol. Briefly spin down the beads and remove the residual ethanol as completely as possible, then air dry the beads for no more than 2 min.
57. Resuspend immediately the beads in 100 µl of water, ready for the next phosphorylation step.

| Fill-in mix (add in this order) | Per reaction |
| --- | --- |
| Bridge adaptor w/ Biotin (40μM) | 25 μl |
| Blunt adaptor w/o Biotin (50 μM) | 20 μl |
| 10x T4 ligase buffer | 10 μl |
| PEG-4000 (50%) | 10 μl |
| T4 DNA ligase (5 U/μl) | 5 μl |

### 3.6. Phosphorylation

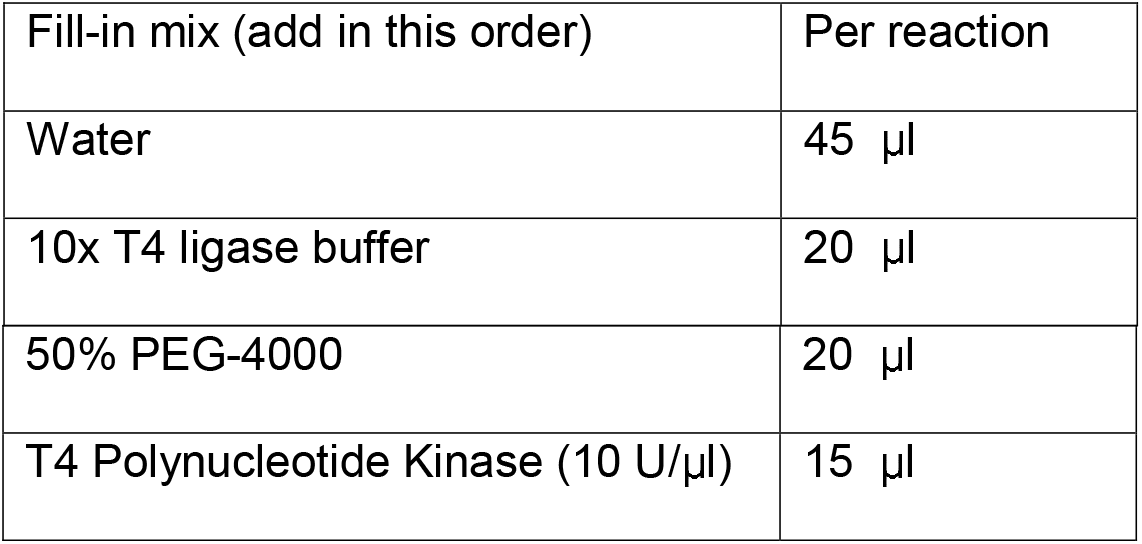
58. Prepare the PNK master mix as follows:
59. Add the 100 µl PNK master mix to the sample of step 57. Mix well by pipetting up and down.
60. Incubate at 37^o^C for 1 h.

### 3.7. In-gel proximity ligation

61. Prepare 1.2% LMP agarose gel stock solution by adding 1.2 g LMP agarose to 100 ml ddH_2_O and melted in a microwave oven.
62. Keep the 1.2% LMP agarose gel stock solution in a 37^o^C water bath.
63. Prepare a 15 ml tube and add 5.7 ml ddH_2_O and 1.1 ml 10x T4 ligase buffer.
64. Transfer the 200 µl of the above phosphorylation reaction mixture of step 60 to the 15 ml-tube.
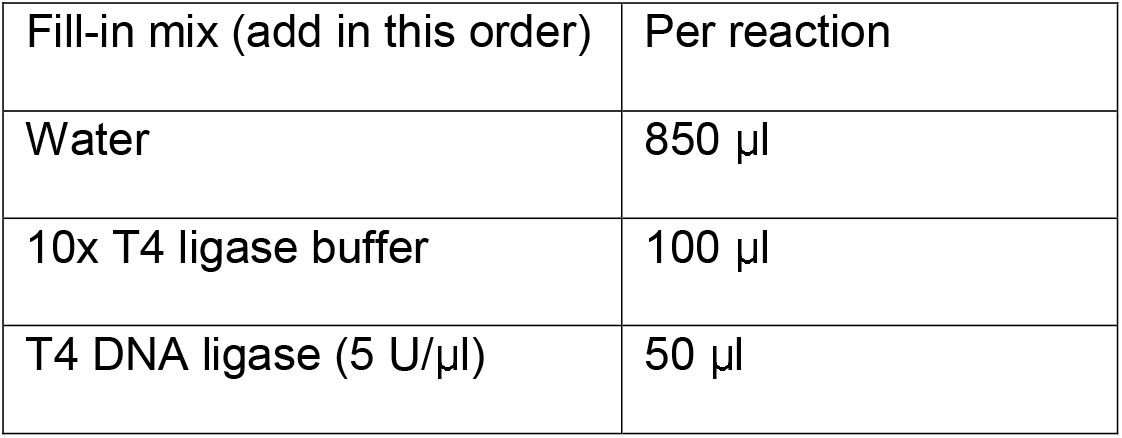
65. Prepare the diluted T4 DNA ligase mix as follows,
66. Add 4 ml of the 1.2% LMP agarose gel stock solution to the 15 ml tube of step 64.
67. Add the 1ml diluted T4 DNA ligase mix to the 15 ml-tube, mix thoroughly immediately by inverting the tubes upside down several times.
68. Immediately place the 15 ml-tube in iced water to solidify the gel.
69. Incubate at 16^o^C for 4 h.
70. Incubate at RT for 1 h.

### 3.8. Reverse cross-linking and DNA purification

71. Incubate the sample from step 70 at 70^o^C for 10 min.
72. Incubate at 42^o^C for 10 min.
73. Add 30 µl of agarase (0.5 u/µl) to the tube.
74. Incubate at 42^o^C overnight.
75. Spin down the magnetic beads in each tube by centrifuging for 1 min at 2987*xg* (Sorvall 75006441 swinging bucket rotor) at RT.
76. Transfer the 12 ml supernatant to an Amicon Ultra-15 Centrifugal Filter Unit (NMWL, 30 KDa, Millipore) and concentrate the volume to about 300 µl by centrifuging for 20 min at 2987*xg* (Sorvall 75006441 swinging bucket rotor) at RT.
77. Transfer both the magnetic beads from step 75 and the 300 µl solution from step 76 to a new 1.7 ml microtube and bring up the volume to 1 ml by adding about 700 µl of 1x TE lysis buffer.
78. Then add100 µl of 20 mg/ml proteinase K to the tube.
79. Incubate overnight at 65^o^C.
80. Add an additional 10 µl 20 mg/ml proteinase K to the tube the next day, and incubate at 55^o^C for another 2 h.
81. Divide the reaction mixture in the tube into two tubes (~600 µl aliquots for each tube).
82. Precipitate DNA with 3 µl GlycoBlue (Ambion), 60 µl 3M Na-acetate (pH 5.2) and 600 µl of iso-propanol at -80^o^C for 2 h.
83. Centrifuge for 30 min at the maxi speed with a microcentrifuge at 4^o^C.
84. Resuspend the DNA pellets in each tube with 100 µl ddH_2_O.
85. Add 600 µl QG buffer of the QIAquick Gel Extraction Kit (Qiagen) to each tube.
86. Incubate at RT for 10-20 min.
87. Purify the DNA with the QIAquick Gel Extraction Kit (Qiagen)
88. Elute the DNA with 100 µl ddH_2_O or EB buffer.
89. Determine the concentration of the recovered DNA with a Nanodrop-1000 spectrophotometer (Thermo Scientific). The typical yield should be around 5 µg per 10^6^ cells.

### 3.9. DNA sonication

90. Shear the DNA (5 µg DNA in 100 µl 1x TE lysis buffer) to a size of 100 – 300 bp using a sonicator. For a Covaris S2 instrument use the following parameters: dDuty cycle: 2%, intensity: 5, cycles per burst: 200, set mode: frequency sweeping, continuous degassing, process time: 20 sec, number of cycles: 5.
91. Transfer the 100 µl sonicated DNA solution to a 1.7 ml microtube and bring the volume to 200 µl by adding ddH_2_O.
92. Add 200 µl of AMPure XP beads to the tube, mix thoroughly by pipetting up and down.
93. Incubate at RT for 5 min, and place the three tubes in a DynaMag-Spin magnet (Life Technologies) for 2 min.
94. Discard the supernatant and wash the beads twice with 1 ml of 80% ethanol. Briefly spin down the beads and remove the residual ethanol as completely as possible, then air dry the beads for 10 min.

### 3.10. End Repair and dA-tailing

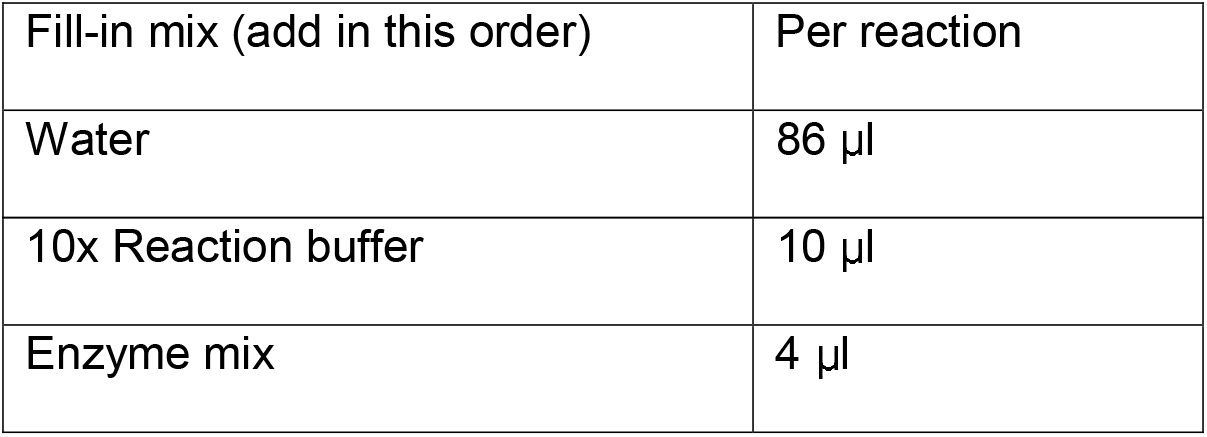
95. Set up the End-repair reaction with the Fast DNA End Repair Kit as follow:
96. Incubate at 16 °C for 10 min.
97. Add 5 µl of 10% SDS to the tube to stop the reaction. Mix thoroughly by pipetting up and down.
98. Add 150 µl of 20% PEG buffer to the tube, mix thoroughly by pipetting up and down.
99. Incubate at RT for 5 min, and place the tube in a DynaMag-Spin magnet for 2 min.
100. Discard the supernatant and wash the beads twice with 1 ml of 80% ethanol. Briefly spin down the beads and remove the residual ethanol as completely as possible, then air dry the beads for 10 min.
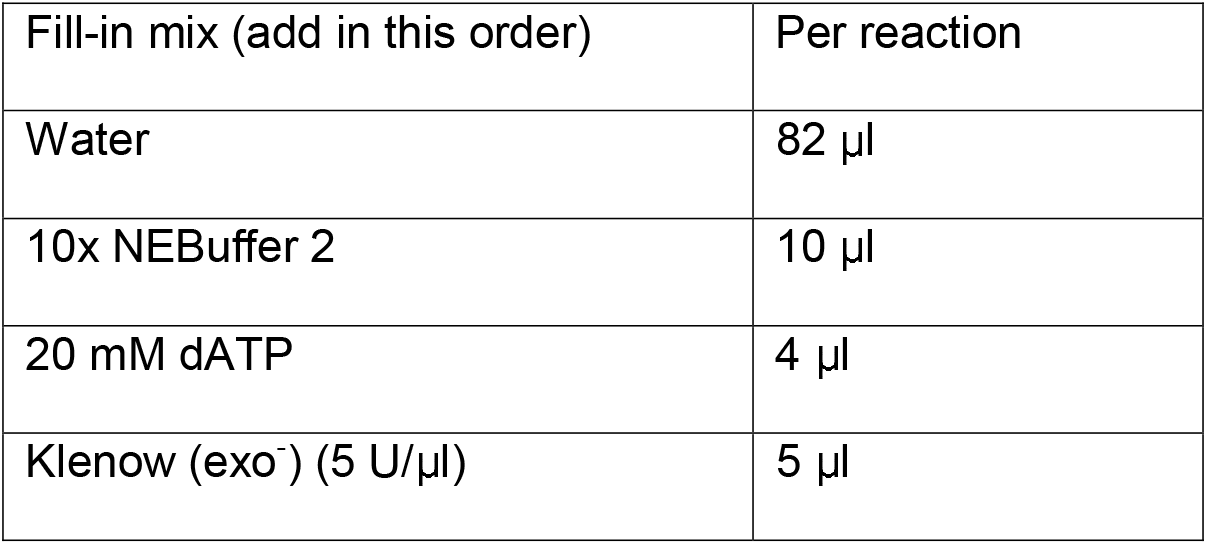
101. Set up the dA-tailing reaction in each tube as follow:
102. Incubate at 37 °C for 30 min.
103. Add 5 µl of 10% SDS to each tube to stop the reaction. Mix thoroughly by pipetting up and down.
104. Add 150 µl of 20% PEG buffer to each tube, mix thoroughly by pipetting up and down.
105. Incubate at RT for 5 min, and place the tubes in a DynaMag-Spin magnet.
106. Discard the supernatant and wash the beads twice with 1 ml of 80% ethanol. Briefly spin down the beads and remove the residual ethanol as completely as possible, then air dry the beads for 10 min.

### 3.11. Ligation of sequencing adaptors

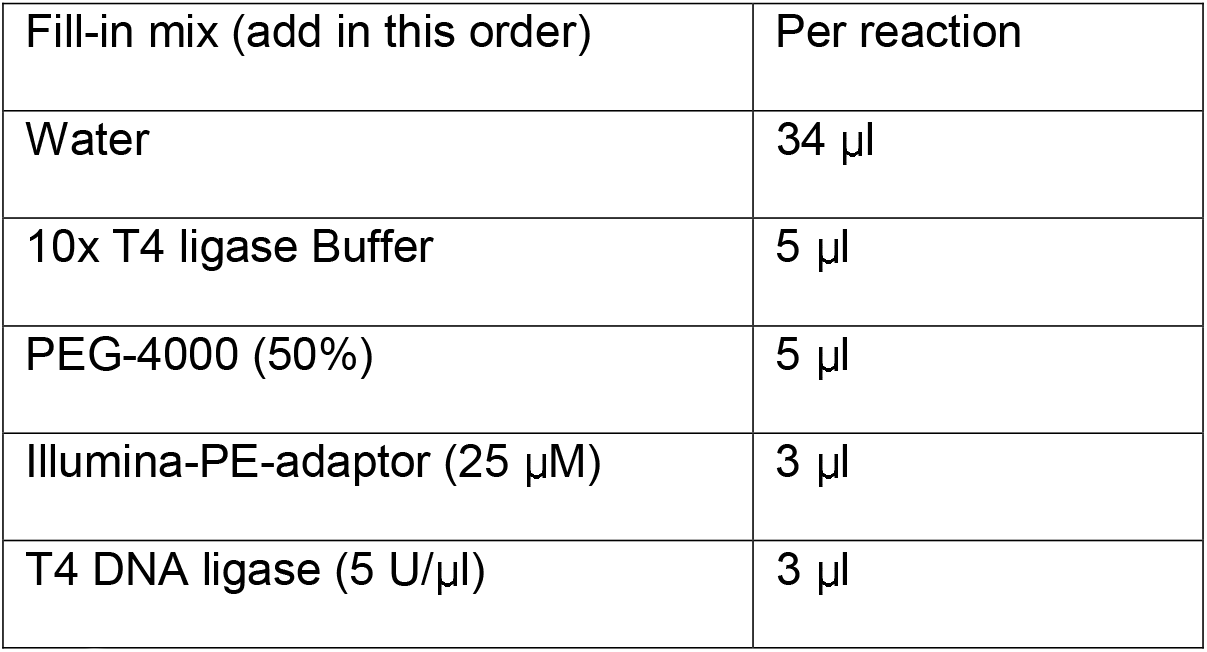
107. Set up the ligation reaction as follow:
108. Incubate at RT for 30 min or 16 °C overnight.
109. Add 5 µl of 10% SDS to each tube to stop the reaction.
110. Add 145 µl of ddH_2_O to bring up the volume to 200 µl, Mix thoroughly by pipetting up and down.
111. Add 200 µl of 20% PEG buffer, mix thoroughly by pipetting up and down.
112. Incubate at RT for 5 min, and place the tube in a DynaMag-Spin magnet for 2 min.
113. Discard the supernatant and wash the beads once with 1.5 ml of 80% ethanol. Briefly spin down the beads and remove the residual ethanol as completely as possible, then air dry the beads for 10 min.
114. Elude DNA from the beads with 100 µl of water, ready for the next biotin pull-down step.

### 3.12. Biotin pull-down

1. 115. Wash 30 µl of MyOne C1 Dynabeads one with 100 µl of 1× B&W buffer, and then resuspend the beads in 100 µl of 2× B&W buffer.
2. 116. Transfer the 100 µl eluted DNA from step 114 to the above tube containing the MyOne C1 beads, mix well.
3. 117. Incubate the sample for 15 min at RT with rotation.
4. 118. Reclaim beads against the DynaMag-Spin magnet for 1 min, discard the supernatant.
5. 119. Wash beads four times with 200 µl of 1X B&W buffer containing 0.1% Tween-20.
6. 120. Wash beads twice with 200 µl of EB buffer.
7. 121. Resuspend the beads in each tube with 40 µl of EB buffer.

### 3.13. Amplification of whole-genome DNase Hi-C library for sequencing

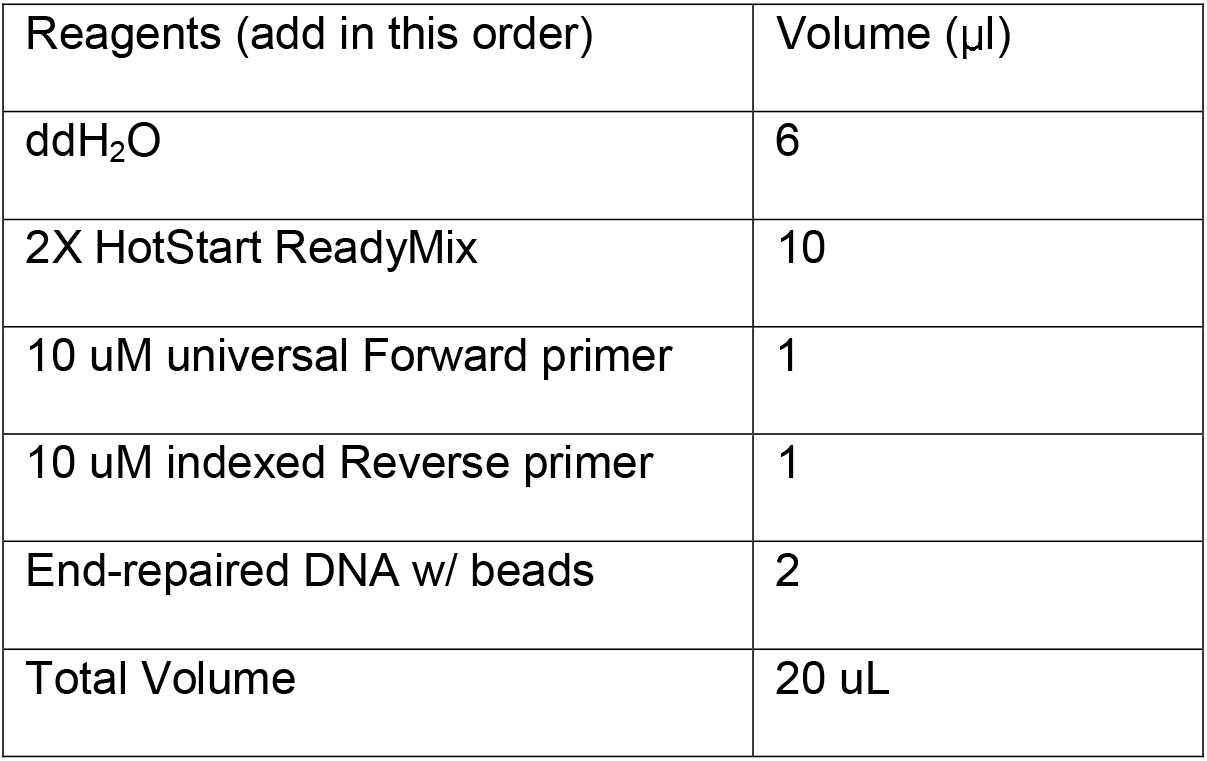
122. To determine the number of PCR cycles necessary to generate enough PCR products for sequencing, set up trial PCR reactions with 8, 10, or 12 cycles, KAPA HiFi HotStart DNA Polymerase and 2 µl of DNA-bound streptavidin beads as follows:
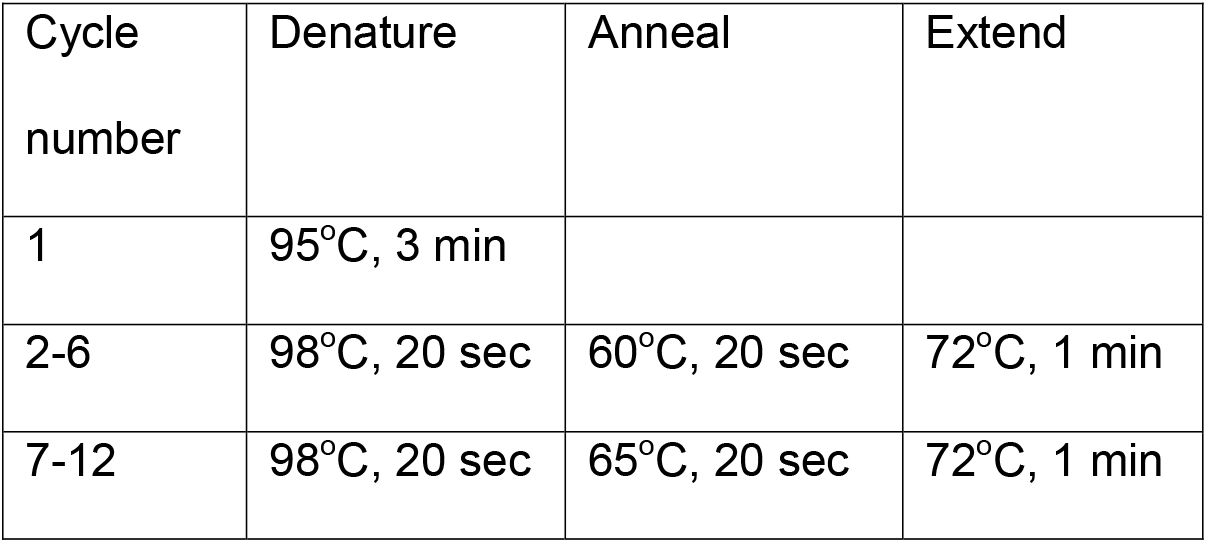
Using the following PCR program:
123. Run out 2 µl of each PCR reaction on a 2% agarose gel to determine the appropriate number of cycles and amount of input beads for each PCR reaction.
124. Aliquot remaining beads into 40 µl PCR reaction and amplify the remaining beads using multiple PCR reactions at the optimized PCR cycle number (usually 9-12).
125. Pool all PCR reactions into one 1.5 mL microcentrifuge tube.
126. Purify library by adding 0.8X volumes of AMPure XP beads.
127. Incubate mixture at RT for 5 min and place tube in a DynaMag magnet for 2 min.
128. Discard the supernatant and wash the beads twice with 1 ml of 80% ethanol. Briefly spin down the beads and remove residual ethanol as completely as possible, then air-dry the beads for 6 min.
129. Elute DNA in 30 µl EB buffer.
130. Determine the DNA concentration using Qubit dsDNA HS kit according to manufacturer’s instructions.

### 3.14. Enrichment of a subset of chromatin interactions of interest using targeted DNase Hi-C

131. To generate DNA templates for targeted DNase Hi-C assays, the above whole-genome chromatin interaction library from step 121 is amplified for 8-9 PCR cycles with KAPA HiFi HotStart DNA Polymerase and the Pre-Capture PCR primer pairs.
132. Design of a bait library to enrich the chromatin interactions associated with specific genomic regions of interest using the design tool of a selected targeted DNA sequencing platform (e.g., the NimbleDesign tool of the SeqCap EZ system (Roche NimbleGen)).
133. Enrich a subset of chromatin interactions of interest using the bait library according to the manufacturer’s instructions. When the SeqCap EZ system is used, about 1 µg of DNA template (from step 13) is required and human Cot-1 DNA is used to block the repetitive regions in the human genome. Also, 1000 nmol of each of the three block oligos, Adaptor-Hi-block, NBGN-8bp-ID-BL, and internal-adaptor-block, are used to block the adaptor sequences in each capture reaction.
134. Amplify and purify the targeted DNase Hi-C library as described in steps 122 to 130. The optimized PCR cycle number is dependent on the size of the targets and the efficiency of the capture reaction.

### 3.15. Quality control of whole genome or targeted DNase Hi-C libraries by BamHI digestion

135. Digest a small aliquot of the final DNase Hi-C library with BamHI to estimate the portion of molecules with valid biotinylated junctions
136. Incubate at 37 °C for 20 min.
137. Run the entire volume of the reaction on a 2% agarose gel (Fig. 3C).

| Reagent | Digestion | No-digestion Control |
| --- | --- | --- |
| 10x Fast digestion buffer | 1 μl | 1 μl |
| DNase Hi-C product | 1-2 μl (5 -10 ng) | 1-2 μl (5-10 ng) |
| Fast digestion BamHI | 1 μl | 0 μl |
| Water | to 10 μl | to 10 μl |

## 4. Data analysis, interpretation and expected results

### 4.1. Mapping, filtering, normalization, and visualization

If DNase Hi-C libraries are sequenced in a multiplexed manner, we first demultiplex the reads based on the corresponding barcodes. Since the chimeric DNA fragments in a DNase Hi-C library harbor both the Illumina primer and the bridge adaptor sequences (Fig. 4A), we then perform an exhaustive search and trimming of these sequences in the full-length reads and extract the remaining read fragments of various lengths (≥25 bp). We then map each end of these cleaned paired-end reads separately to a reference genome and only retain the read pairs with both reads mapped uniquely, allowing at most 3 mismatches and requiring mapping quality MAPQ≥30. Finally, PCR duplicates are removed by eliminating redundant paired-end reads, defined as those pairs where both ends were mapped to identical locations in the genome.

We use the resulting valid read pairs to generate whole-genome contact matrices at both 1 Mb and 40 kb resolutions (the resolution is variable depending on the sequencing depth). To do so, we partition the whole genome into non-overlapping bins and count the number of contacts (i.e., mapped paired-end reads) observed between each pair of bins. The dimension of the resulting contact map is the total number of bins in the genome, and entry (i, j) is the contact count between bins i and j. We then normalize the whole-genome contact map using an iterative correction method (ICE) [53] to obtain a normalized contact map with near-equal row and column sums. Prior to applying the iterative correction procedure, the contact maps are filtered as follows: bins along the diagonal, super-diagonal (+1 off-diagonal) and sub-diagonal (-1 off-diagonal) (representing entries dominated by unligated fragments (dangling ends) and self-ligation products) and bins with the lowest 2% read coverage (representing unmappable regions and sparsely populated regions dominated by spurious contacts) have their contact counts set to zero. Additionally, to facilitate the comparison of contact maps across multiple samples, intra-chromosomal contact maps are scaled to the same sequencing coverage.

In addition to the pipeline described here, there are a number of public pipelines for processing Hi-C data (reviewed in [54]), as well as tools for visualizing Hi-C contact matrices (reviewed in

[55]). These pipelines and tools can also be adapted to whole-genome DNase Hi-C datasets.

### 4.2. Identifying architectural features of 3D genome organization using whole-genome DNase Hi-C datasets

Whole-genome DNase Hi-C data can be used to identify the multiple layers of architectural features of the hierarchical 3D genome organization, such as chromosome territories, A/B compartments, TADs and chromatin loops. Chromosome territories can be visualized by generating a whole-genome contact map using the inter-chromosomal interactions between each chromosome pair of a genome [20] [56]. A/B compartments can readily be identified by using principal component analysis (PCA), since these compartments tend to be captured by the first principal component [20]. TADs can be mapped when interaction data is binned at 40 kb or less, using any one of a number of available algorithms (reviewed in [57] and [58]). Unlike chromosome territories, A/B compartments, or TADs, whose identification requires only moderate Hi-C data resolutions, chromatin loops can only be identified at much higher resolution (e.g., a 10 kb or higher resolution) [27]. However, even at lower resolution, it is possible to identify statistically significant contacts, subject to, for example, a 1% false discovery rate threshold, by using a straightforward statistical model [59].

DNase Hi-C can also identify other conformational features of chromosome folding. For example, we have applied DNase Hi-C combined with allele-specific analyses to determine the 3D architecture of the X chromosomes, and obtained the first 3D view of the whole inactive X chromosome, thereby revealing a surprising bipartite structure with an apparent hinge that binds CTCF on the inactive allele [46].

### 4.3. Identifying the landscapes of physical interactomes between cis-regulatory elements using targeted DNase Hi-C datasets

Targeted DNase Hi-C is designed for fine-mapping chromatin architecture of specific genomic regions of interest at unprecedented resolution in a massively parallel fashion, which is well suited for high-throughput mapping (surveying tens of thousands of targets in a single experiment) of fine-scale 3D architecture, and in particular, the 3D organization of *cis*-regulatory elements [44]. Here we outline the basic data analysis pipeline for processing targeted DNase Hi-C datasets.

#### 4.3.1. Identification of captured target-associated chromatin interactions

We trim, map and filter the paired-end sequencing reads from targeted DNase Hi-C data in a similar fashion to that of the whole-genome DNase Hi-C data as described in Section 4.1. However, for targeted DNase Hi-C data we perform an additional filtering step to keep only the paired-end reads for which at least one end mapped within 150 bp of one of the target regions. We then use these target-associated reads to define contact maps of the captured target regions at 1 kb and 10 kb resolutions (depending on sequencing depth).

#### 4.3.2. Assessing the enrichment efficiency of each target

We next measure the capture efficiency of each target region as its captured read coverage (number of captured read pairs per kilobase of the target region length). We rank the targets based on their capture efficiency and exclude those targets with very low captured read coverage (lower than 25% of the average captured read coverage among all targets) in further analyses. These low-coverage targets are mainly located in unmappable genomic regions.

#### 4.3.3. Normalization of targeted DNase Hi-C data

To correct biases in targeted DNase Hi-C data, we estimate a bias factor for each bin (either 1 kb or 10 kb). To do so, we first set the bias of unmappable bins (mappability score < 0.5) to be 1. A mappability score less than 0.5 for a bin means that more than half of the bases in that bin are not uniquely mappable for 50-bp reads. We calculate mappability scores using GEM [60]. Then we assess the biases in target bins (those overlapping with the designed target regions on the DNA capture array) and non-target bins, separately, using the following strategies.

For bins overlapping with target regions, we approximate their biases by measuring their coverage in the targeted DNase Hi-C data. The bias factor is calculated as the number of captured read pairs at each target region per kilobase of the target region length. Bins overlapping with the same target region share the same bias factor. We then normalized the bias factors at target bins so that their average equals 1.

For bins that do not overlap with any target, we estimate their biases from a corresponding parent whole-genome DNase Hi-C library. First, assuming, as in the ICE method [53], that all non-target bins have equal “visibility”, we take the bin coverage (i.e., the row margins of the contact matrix) at either 1 kb or 10 kb resolution, normalized by dividing by the average among all mappable bins in the genome. We then truncate the normalized bin coverage at 5% and 95% percentiles and perform smoothing by taking the average of 10 neighboring bins. Our normalization method is similar to ICE, in the sense that taking the contact margins is equivalent to the ICE correction with only one iteration. Accordingly, we have observed that contact margins and ICE iteratively learned bias factors are highly correlated at 40 kb resolution. In our case, we chose not to perform iterative corrections at 1 kb resolution because the contact matrix becomes very large and sparse at 1 kb resolution. Consequently, the normalization procedure is computationally expensive and is also unstable.

#### 4.3.4. Identifying statistically significant contacts associated with each target

Statistically significant contacts associated with the target regions can be identified using the Fit-Hi-C method [59]. To apply the Fit-Hi-C method to targeted DNase Hi-C datasets, we made two modifications to the original method. First, because in targeted DNase Hi-C experiments, DNase

I was used instead of restriction enzymes, we aggregated chromatin contacts using fixed sized bins (either 1 kb or 10 kb) instead of aggregating within restriction enzyme fragments. Second, we estimated the null contact probability using only the pairs of loci with at least one end overlapping one of the captured target regions.

For short-range (< 10 Mb) intra-chromosomal chromatin contacts, we first parse the target-captured reads at 1 kb resolution and then apply the modified Fit-Hi-C method to estimate the null distribution of contacts within the genomic distance range of 5 kb to 10 Mb. We discard the very short-range contacts (< 5 kb) because they are mainly self-ligation products or dangling ends. We use one round of refinement (i.e., two rounds of spline-fitting) to estimate this null distribution and then identify the significant contacts at false discovery rate (FDR) < 0.05. For long-range (≥ 10Mb) intra-chromosomal chromatin contacts, we parse the target-captured reads at 10 kb resolution and then use the modified Fit-Hi-C method to estimate the null distribution of contacts within the full range of full chromosome length using one round of refinement. Similar to short-range contacts, we identify significant contacts at FDR < 0.05. For inter-chromosomal contacts, we identify significant target-captured contacts at 10 kb resolution using a simple binomial model, as described in [56] and use a FDR < 0.05. To eliminate contacts that are introduced by mapping biases at low-mappability regions, we further discard contacts that are associated with genomic bins that have low mappability score (< 0.5).

In the event of a bona fide chromatin looping contact, we expect the immediately flanking bins around the contacting regions to be also within relatively close proximity [44]. Thus, we further apply a neighborhood filter using the contact significance we computed from Fit-Hi-C to identify highly confident contacts associated with each target, as follows. For each chromatin contact between target t and non-target genomic bin i, we call it a high confidence contact if the contact itself meets the stringent FDR cutoff of 0.05, and at least 3 out of 10 neighboring bins (5 on each side) of bin i contact with the target t at a permissive FDR cutoff of 0.1. After the neighborhood filtering, we merge adjacent and nearby (< 3 bins apart) high confidence contact bins associated with the same target.

#### 4.3.5. Integrative analysis with functional genomic features

Integrative analysis can be carried out by overlaying targeted DNase Hi-C datasets with other genomic, epigenomic and transcriptomic datasets, including gene expression profiles, chromatin states, chromatin accessibility landscapes, enhancer and super-enhancer annotations and disease-associated SNPs. The unprecedented resolution of targeted DNase Hi-C is approaching that of annotations of functional genomic *cis*-elements, thereby allowing for the detailed integration of 3D genome architecture and one-dimensional epigenomic information. We found that promoter-associated chromatin contacts are significantly enriched with *cis*-regulatory elements such as enhancers, super-enhancers and transcription factor binding sites. Hence, targeted DNase Hi-C can be used to delineate the *cis*-regulatory networks that control gene expression. One of the straightforward applications of targeted DNase Hi-C will be to systematically link disease-associated non-coding SNPs to their target genes in the context of nuclear 3D organization. Targeted DNase Hi-C may also prove to be valuable for characterizing phenotype-associated chromatin 3D signatures and for probing the relationship between genome architectural defects and disease pathogenesis.

#### 4.3.6. Visualization of targeted DNase Hi-C contact profiles

Intra– and inter-chromosomal contact profiles of each target can be visualized with domainograms using the 4Cseqpipe software [61] and Circos diagrams [62], respectively (Fig. 5).

**Fig. 5.**
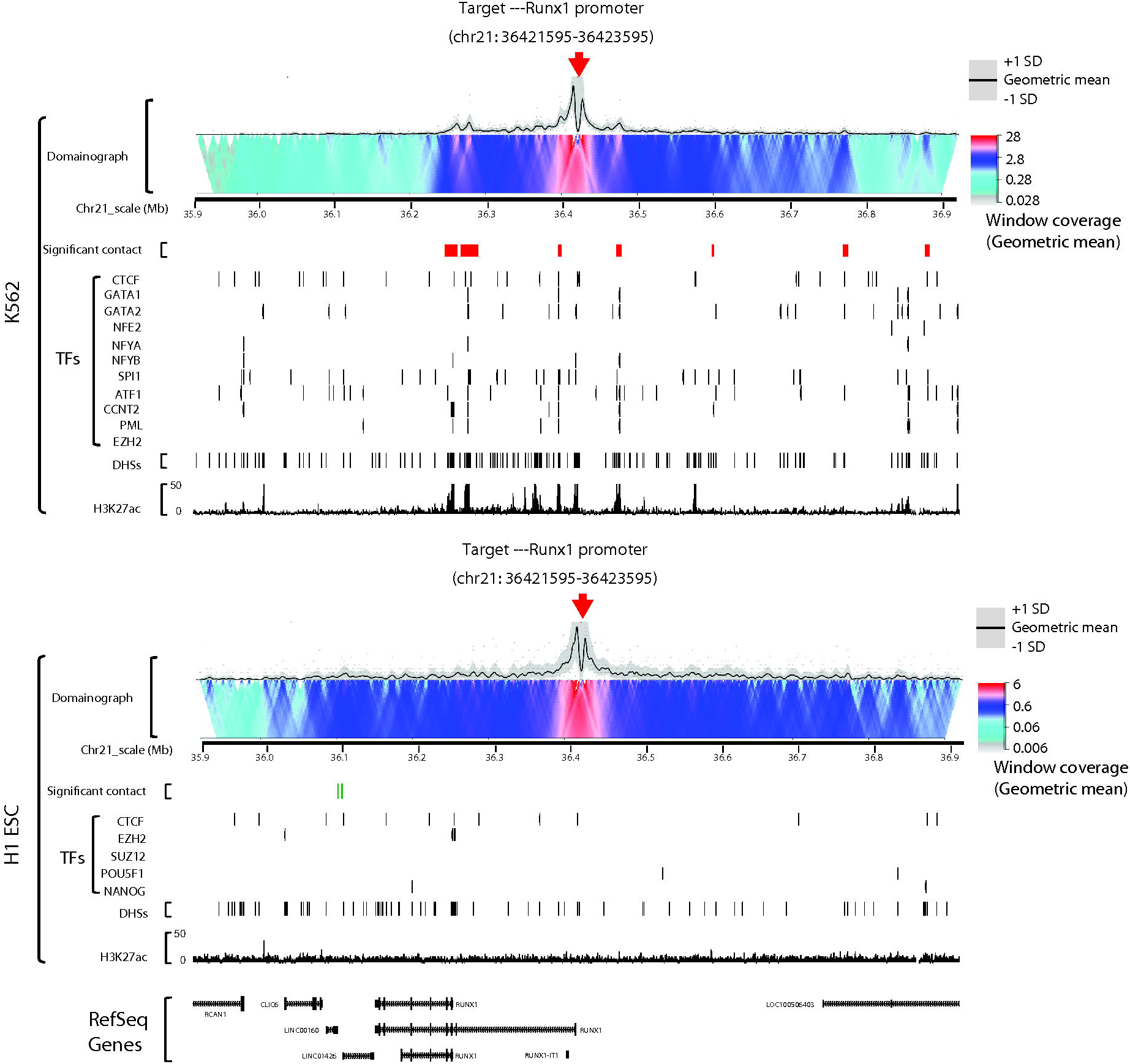
Visualization of targeted DNase Hi-C results using Circos diagram and domainograms. (A) Circos plots showing representative targeted DNase Hi-C-identify high confidence cell-type specific inter-chromosomal contacts in H1 and K562 cells. Each arc represents a contact; different colors are used to distinguish individual chromosomes. Targets are HS2-HS3 on chr11, Nanog on chr12, and Sox2 on chr3. (B) Cell type-specific 3D organization of the RUNX1 promoter in K562 and H1 cells. Domainographs showing contact profiles and significant contacts (red or green boxes) within 500 kb (upstream or downstream, chr21:35,921,595-36,923,595) of the target (RUNX1 promoter) are shown. Red arrows indicate the position of the target. The tracks of H3K27ac and DNase hypersensitive sites (DHSs) for both K562 and H1 ESCs are snapshots taken from the UCSC Genome Browser using data from the ENCODE Project Consortium. UCSC Genome Browser tracks for TFBS (CTCF, GATA-1, GATA-2, NF-E2, NF-YA, NF-YB, SPI1, ATF1, CCNT2, PML and EZH2 for K562cells; CTCF, EZH2, SUZ12, POU5F1, and NANOG for H1 ESCs) are also shown. The RefSeq genes are shown in the bottom. Chromosomal position is based on GRCh37/hg19.

## 5. Conclusions

DNase Hi-C protocols employ DNase I digestion to fragment chromatins, providing a method with unlimited methodological resolution for mapping 3D genome architecture.

## 6. Appendixes

### Buffers and Materials

#### Buffers

##### 1X Cell Lysis buffer

10 mM Tris-HCl pH8.0

10 mM NaCl

0.2% Igepal CA-630 (NP40)

Autoclaved water

— Store at 4°C

##### 1x TE lysis buffer

50mM Tris.HCl (pH7.0)

1mM EDTA

1% SDS

— Store at RT

##### 5x stop buffer

125 mM EDTA

3% SDS

— Store at RT

##### 20% PEG buffer

20% (v/v) PEG-8000

2.5 M NaCl

##### 2X B & W buffer

10 mM Tris-HCl pH8.0

1 mM EDTA

2 M NaCl

##### 1XB & W buffer with 0.1% Tween-20

5 mM Tris-HCl pH8.0

0.5 mM EDTA

1 M NaCl

0.1% (v/v) Tween-20

##### EB buffer

10 mM Tris-HCl pH 8.5

## Acknowledgements

This work was supported by the NIH Common Fund U54DK107979 to XD, CMD, WSN, ZD and JS) and UW Bridge Fund to ZD. JS is an Investigator of the Howard Hughes Medical Institute.

